# Chronic Benzene Exposure Aggravates Pressure Overload-Induced Cardiac Dysfunction

**DOI:** 10.1101/2021.08.31.458367

**Authors:** Igor N. Zelko, Sujith Dassanayaka, Marina V. Malovichko, Caitlin M. Howard, Lauren F. Garrett, Uchida Shizuka, Kenneth R. Brittian, Daniel J. Conklin, Steven P. Jones, Sanjay Srivastava

**Author notes:** Address correspondence to: Igor N. Zelko, PhD, Department of Medicine, Division of Environmental Medicine, Room 310 CII Building, 302 E. Muhammad Ali Blvd, Louisville, KY 40202, or Sanjay Srivastava, PhD, Superfund Research Center, Room 306 CII Building, 302 E. Muhammad Ali Blvd, Louisville, KY 40202.

## Abstract

Benzene is a ubiquitous environmental pollutant abundant in household products, petrochemicals and cigarette smoke. Benzene is a well-known carcinogen in humans and experimental animals; however, little is known about the cardiovascular toxicity of benzene. Recent population-based studies indicate that benzene exposure is associated with an increased risk for heart failure. Nonetheless, it is unclear whether benzene exposure is sufficient to induce and/or exacerbate heart failure. We examined the effects of benzene (50 ppm, 6 h/day, 5 days/week, 6 weeks) or HEPA-filtered air exposure on transverse aortic constriction (TAC)-induced pressure overload in male C57BL/6J mice. Our data show that benzene exposure had no effect on cardiac function in the Sham group; however, it significantly compromised cardiac function as depicted by a significant decrease in fractional shortening and ejection fraction, as compared with TAC/Air-exposed mice. RNA-seq analysis of the cardiac tissue from the TAC/benzene-exposed mice showed a significant increase in several genes associated with adhesion molecules, cell-cell adhesion, inflammation, and stress response. In particular, neutrophils were implicated in our unbiased analyses. Indeed, immunofluorescence studies showed that TAC/benzene exposure promotes infiltration of CD11b^+^/S100A8^+^/myeloperoxidase^+^-positive neutrophils in the hearts by 3-fold. *In vitro*, the benzene metabolites, hydroquinone and catechol, induced the expression of P-selectin in cardiac microvascular endothelial cells by 5-fold and increased the adhesion of neutrophils to these endothelial cells by 1.5-2.0-fold. Benzene metabolite-induced adhesion of neutrophils to the endothelial cells was attenuated by anti-P-selectin antibody. Together, these data suggest that benzene exacerbates heart failure by promoting endothelial activation and neutrophil recruitment.

## Introduction

Environmental pollution is a health problem worldwide. More than 9 million premature deaths are attributed to pollution every year, out of which 6 million deaths are related to air pollution (1). Nearly half of the air pollution-associated deaths are ascribed to cardiovascular disease (1). Cardiovascular health effects of criteria pollutants (particulate matter2.5, sulfur dioxide, carbon monoxide, nitrogen oxides, ozone and lead) have been extensively studied in the last three decades; however, information about the potential cardiovascular toxicity of other airborne chemicals, such as volatile organic compounds (e.g. benzene, 1,3-butadiene, vinyl chloride, trichloroethylene etc.), is sparse. Benzene, a well-known carcinogen, is abundant in household products, cigarette smoke and automobile exhaust (2–5), and the atmospheric concentration of benzene can exceed 50 ppm, especially near the emission source. People working at gasoline pumping stations or living near hazardous waste sites can be exposed to high levels of benzene. The United States Occupational Safety and Health Administration has set the occupational benzene exposure limit of 1 ppm (6); however, occupational benzene exposure in excess of 100 ppm has been reported in developing countries (7).

Like pollution, heart failure is a pervasive health problem. Heart failure affects more than 6.5 million people in the United States (8) and 26 million people worldwide (9). The lifetime risk for heart failure at 40 years of age is estimated to be 1 in 5 people (10). The pathogenesis of heart failure is also not clearly understood, although a substantial body of literature points to the important role of infiltrating immune cells in the development of left ventricular hypertrophy and cardiac dysfunction (11–18). Therefore, the current focus in finding new therapeutic approaches to delay advanced heart failure includes understanding the molecular mechanisms that govern cardiomyocyte interactions with immune cells (19–21). The innate immune response induced by heart failure initiates the cardiac repair process, and involves infiltration of peripheral neutrophils (22–24).

Recent population based studies have reported that benzene exposure is associated with an increased risk for heart failure (25), and presence of mono-nitrogen oxide and benzene in the air of hospital wards is associated with a higher risk of heart failure morbidity (26). Other studies have linked exposure to benzene with cardiovascular mortality (27, 28); however, direct effects of benzene exposure on heart failure in well-controlled animal studies have not been studied. Here, we examined the effect of chronic benzene exposure on transverse aortic constriction (TAC)-induced cardiac function and associated immune response in mice.

## MATERIALS AND METHODS

### Reagents

Benzene permeation tubes were obtained from Kin-Tek (La Marque, TX). Primers and probes for real-time PCR were purchased from Integrated DNA Technologies (Coralville, IA) and ThermoFisher Scientific (Waltham, MA). Sources of antibodies used for immunohistochemistry and functional studies were: anti-CD11b, myeloperoxidase (Abcam); anti-S100A8 (63N13G5)-FITC from Novus (Novus, Centennial, CO); anti-CD90/Thy1 from Sino Biological (Sino Biological); CD62P from Biolegend (San Diego, CA). Murine cardiac microvascular endothelial cells (CMVEC) were obtained from CellBiologics, Chicago, IL. All other chemicals and enzymes were from Sigma Chemical Co. (St. Louis, MO), or Invitrogen (Carlsbad, CA).

### Animal housing and maintenance

C57BL/6J male mice obtained from Jackson Laboratory (Bar Harbor, ME) were maintained on normal chow in a pathogen-free facility accredited by the Association for Assessment and Accreditation of Laboratory Animal Care. All procedures were approved by the University of Louisville Institutional Animal Care and Use Committee.

### Animal Surgeries

For TAC (29), male C57BL/6J mice 12 weeks of age were anesthetized (intraperitoneal injections of 50 mg/kg sodium pentobarbital and 50 mg/kg ketamine hydrochloride), antiseptically prepared for surgery, orally intubated, and ventilated (oxygen supplement to the room-air inlet) with a mouse ventilator (Hugo Sachs). Core body temperature was maintained at 36.5–37.5°C with an automatic, electronically regulated heat lamp. The aorta was visualized following an intercostal incision. A 7-0 nylon suture was looped around the aorta between the brachiocephalic and left common carotid arteries. The suture was tied around a 27-gauge needle placed adjacent to the aorta to constrict the aorta to a reproducible diameter. The needle was removed, and the chest was closed in layers. Mice were extubated upon recovery of spontaneous breathing. Analgesia (ketoprofen, 5 mg/kg) was provided prior to recovery and by 24 and 48 h post-surgery. Sham mice were subjected to the same procedure as the TAC cohort except the suture was not tied.

### Benzene Exposure

One week after the TAC, mice were exposed to benzene for 6 weeks in bedding-free cages as described before (30). Briefly, benzene atmospheres were generated from liquid benzene (Sigma-Aldrich) in a KIN-TEK Analytical, Inc permeation tube. A carrier gas (N2) was delivered to the permeation tube at 100 ml/min and diluted with HEPA- and charcoal-filtered room air (3 L/min) and diluted gas directed to an exposure unit. Flow was distributed through a fine mesh screen of a custom cyclone-type top (Teague Enterprises) that distributed air within 10% of the mean concentration at six locations in the cage. Throughout the exposure, benzene concentrations were continuously monitored using an in-line photoionization detector (ppb RAE: Rae Industries) upstream of the exposure unit. Mice were exposed to 50 ppm benzene (6 h/day, 5 days/week) for 6 weeks. Mice exposed to HEPA- and charcoal-filtered room air only were used as controls.

### Echocardiography

At the end of the benzene exposure protocol, mice were anesthetized with 2% isoflurane, and echocardiography was performed (VisualSonics Vevo 3100), similar to our previous reports (29, 31). At the end of the experiment, hearts were excised for biochemical and pathological analyses. For mice in the TAC arm of the study, Doppler verification of the stenosis was used. The sonographer was blinded to the specific experimental group.

### RNA Isolation and RNAseq analysis

Total RNA was extracted from the hearts of mice using TRIzol kit (Thermo Fisher Scientific, MA, USA), and the purity of RNA was analyzed using NanoDrop One (ThermoScientific, MA, USA). RNA quality was measured by Agilent 2100 bioanalyzer (Thermo Fisher Scientific, MA, USA) and samples with high RNA integrity were used for subsequent RNAseq analysis. RNA samples were processed by Novogene using mRNA and small RNA sequencing services (Novogene, Beijing, China). The resultant raw reads of the FASTQ files were processed and quality metrics were visualized using FastQC v 0.11.9. The mRNA differentially regulated genes (DEG) and pathway enrichment analysis were performed using the NGS Data Analysis pipeline (32).

### Histopathology

Formalin-fixed, paraffin-embedded hearts from Sham and TAC mice were sectioned at 4 μm thickness, and later deparaffinized, and rehydrated according to the appropriate staining method. The histological staining was performed as previously described (33, 34). Picro-Sirius Red staining was used to evaluate tissue fibrosis and general histology, and Alexa Fluor 555-conjugated wheat-germ agglutinin (Invitrogen) staining was used to determine the average myocyte area. All images were captured at 20X magnification using the Keyence BZ-X810, all in one fluorescent microscope. Both fibrosis and myocyte area were analyzed using the hybrid cell count feature (BZ-H4C) associated with the Keyence system. Fibrosis was expressed as a percentage of scar tissue divided by the total area of tissue.

### Immunofluorescence

For immunostaining, deparaffinized and rehydrated sections were incubated for 20 min with 10 mmol/L citric acid pH 6.0 (pH 9.0 for MPO staining). Nonspecific binding was blocked with 5% normal goat serum and 0.05% saponin (Sigma) in PBS (pH 7.4) for 30 min. Sections were blocked and then incubated with the appropriate primary antibody in PBS with 1% BSA and 0.05% saponin for 1 h at 37°C against: CD11b (Abcam), S100A8-FITC (Novus), myeloperoxidase (Abcam). Tissue sections were then incubated for 30 min at room temperature with respective secondary antibodies conjugated with Alexa 488, 555, or 647 (Invitrogen), and counterstained with DAPI to label nuclei. Images were made with a 20x and 60x objective.

### Neutrophils isolation

Neutrophils were isolated from bone marrow as described previously (35). Briefly, C57BL6/J male mice 10 weeks old were sacrificed using pentobarbital injection. Bone marrow cells were flushed from the femur and tibia and filtered through a 70 μm mesh filter. For a negative selection of neutrophils, we used a Neutrophil Isolation kit from Miltenyi, Auburn, CA. Magnetic sorting was performed using MS columns. Neutrophils presented in flow-through fractions were characterized using flow cytometry and by cytospin with Diff-Quick staining. Neutrophil purity was always higher than 93%.

### Cell adhesion assay

Neutrophil adhesion to CMVEC was determined as previously described (36). Briefly, CMVEC (passage 6 to 10) were seeded into 96-well plate at 25,000 cells per well in 100 μl of growth media. Twenty-four hours later confluent monolayer of endothelial cells was exposed to benzene metabolite catechol (5 μM) for 24 hours. Cells treated with TNFα (100 ng/mL) for 4 hours served as positive controls. Isolated neutrophils labeled with calcein (36) were added to endothelial cells and incubated for 30 min at 37 °C. Cells were rinsed with PBS containing Mg^2+^ and Ca^2+^ and the fluorescence was measured with a Synergy H1 (Biotek) fluorescence plate reader (excitation wavelength, 485 nm and emission wavelength, 520 nm). To measure the effect of p-selectin blocking on neutrophil adhesion, CMVEC were incubated with anti-CD62P antibodies (25 μg/mL) for 30 min prior to the neutrophil adhesion assay. For each assay, 10-12 wells were used.

### Statistical Analyses

Data are presented as means ± SEM. The statistical significance of differences was determined by t-test. A one-way analysis of variance (ANOVA) with Holm-Sidak *post hoc* test was used to compare differences between multiple treatment groups. P-value of <0.05 indicated statistically significant differences. All analyses were performed using Excel and GraphPad Prism software (GraphPad Software, San Diego, CA). Statistical analyses for echo-based parameters were performed after adjustment for body weight and heart rate by two-way ANOVA with Tukey-Kramer corrections.

## RESULTS

### Benzene exposure exacerbates pressure overload-induced cardiac dysfunction

As expected, mice in the TAC/Air group developed significant left ventricular dilation as indicated by a significant increase in end diastolic volume (EDV), end systolic volume (ESV) and left ventricular internal diameter in diastole (LVIDd), and diameter in systole (LVIDs), and consequent decrease in fractional shortening (FS) and ejection fraction (EF), as compared with the mice in corresponding Sham/Air (**Fig. 1**). TAC also increased the heart mass, cardiac myocyte size and fibrosis in the heart (**Fig. 1**). This was accompanied with a significant increase in the fluid accumulation in the lung (**Supplemental Fig. 1**). Benzene exposure had no effect on myocyte size and cardiac function in the Sham group; however, it significantly compromised cardiac function as depicted by a significant increase in EDV, ESV, LVIDd, and LVIDs and resultant decrease in FS and EF, as compared with TAC/Air-exposed mice (**Fig. 1** and ***Supplemental* Table 1**). The other cardiac function parameters such as left ventricular anterior wall thickness at diastole or systole (LVAWd and LVAWs) and left ventricular posterior wall thickness at diastole or systole (LVPWd and LVPWs), isovolumic relaxation time (IVRT), Stroke Volume (SV), and Cardiac Output (CO) did not significantly change in TAC/Benzene vs TAC/Air groups (***Supplemental* Table 1**). Cardiac myocyte size and fibrosis in TAC/Benzene-exposed mice were also comparable with the TAC/Air-exposed mice. Collectively, these data suggest that a 6-week benzene exposure exacerbates TAC-induced cardiac dysfunction but does not induce cardiac remodeling in Sham mice.

**Figure 1.**
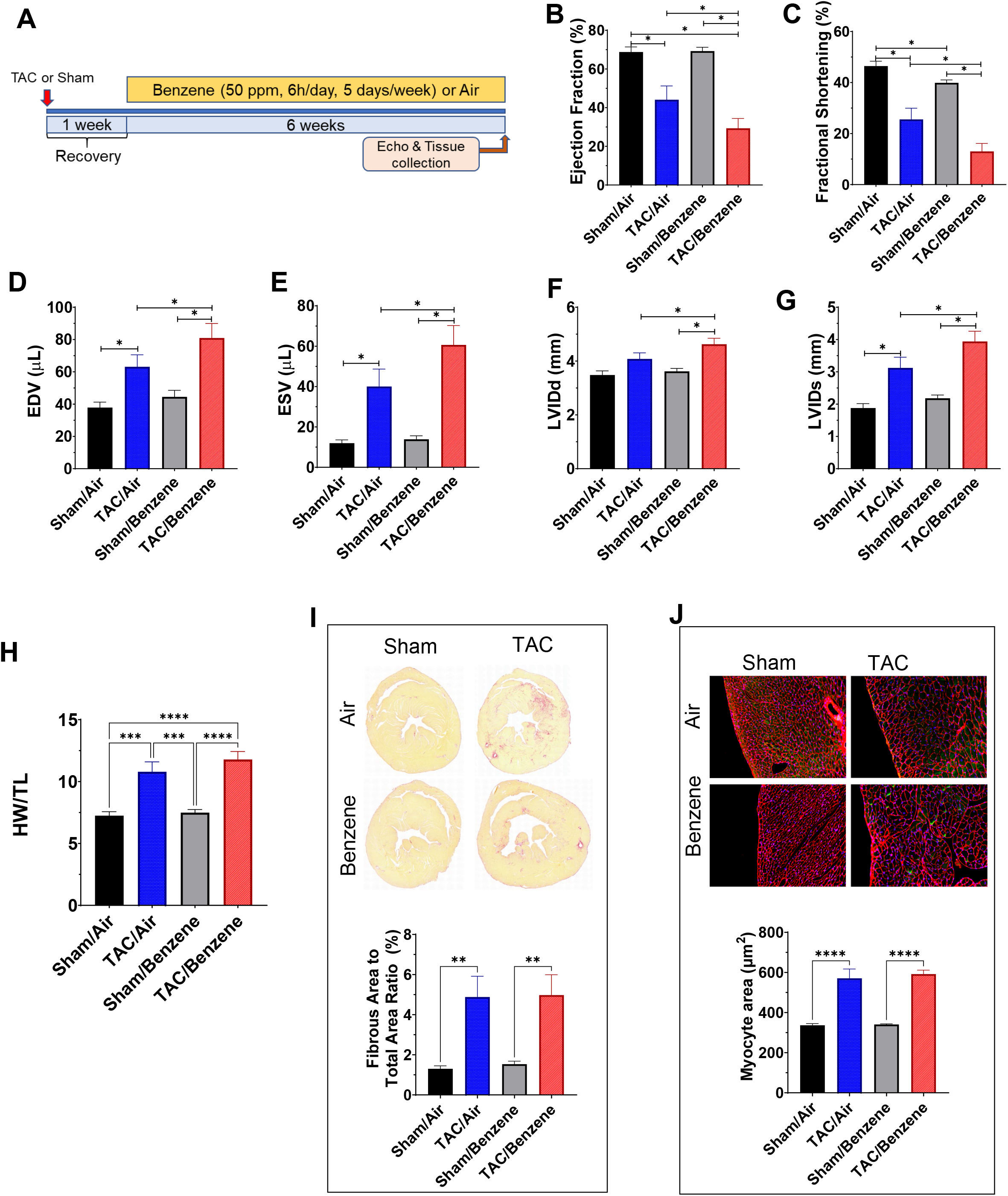
Effect of benzene exposure on left ventricle function and cardiac remodeling after pressure overload. **A.** Benzene exposure protocol: Transverse aortic constriction or Sham operated male C57BL/6 mice were exposed to benzene (50 ppm, 6 h/day, 5 days/week) or HEPA-filtered air (Air) for 6 weeks. At the end of the exposure protocol, echocardiography was used to measure ejection fraction (**B**), fractional shortening (**C**), end-diastolic volume (EDV; **D**), end-systolic volume (ESV; **E**), left ventricular internal diameter end diastole (LVIDd; **F**) and end systole (LVIDs; **G**). Heart weight:tibial length (TL) are presented in panel **H**. Panel **I** shows the representative images of Sirius Red-stained mid-ventricular histological sections from mouse hearts with quantitative analysis of fibrosis expressed as percent of fibrous area to total area ratio. Representative images of wheat germ agglutinin (WGA)-stained mid-ventricular histological sections from mouse hearts with quantitative analysis of myocyte area are illustrated in panel **J**. Values are mean ± SEM. *p<0.05, **p<0.01, ***p<0.001, and ****p<0.0001 represent statistical significance between corresponding groups analyzed by One-way ANOVA. N=6-11/group.

### RNA-Seq analyses of TAC-benzene-exposed hearts

To examine the molecular mechanisms by which benzene worsens cardiac function following TAC, we performed deep RNA sequencing on the left ventricle and septum tissues. For these analyses, we used FDR ≤ 0.01 and 0.5 ≤ Log2 (fold change) ≤ −0.5 to examine the differential expression of the genes. As shown in **Fig 2A**, TAC upregulated 671 genes (e.g., genes involved in extracellular matrix organization and collagen formation) and downregulated 351 genes (e.g., genes involved in muscle contraction and cardiac conduction) in the air-exposed mice (see ***Supplemental Table 1***). Exposure to benzene in Sham-operated mice did not induce any genes while uteroglobin (*Scgb1a1*) and surfactant protein C (*Sftpc*) were downregulated by 8.9- and 6.1-fold. However, benzene exposure in TAC mice significantly upregulated 876 genes and downregulated 674 genes as compared with the corresponding Sham-operated controls. Comparison of the gene expression between TAC/Benzene versus TAC/Air mice showed upregulation of 50 and downregulation of 69 genes. **Fig. 2B** illustrates the heat-map of top the 15 upregulated and 10 downregulated genes between these groups as well as the differential expression of these genes in the other experimental groups. Gene ontology of biological process and molecular function enrichment analyses of differentially expressed genes in the benzene-exposed TAC hearts show significant enrichment of the genes associated with the regulation of “neutrophil degranulation”, “signaling by interleukins”, “extracellular matrix organization”, “platelet degranulation”, and heat shock factor-1 (HSF-1) - dependent transactivation etc. (**Fig. 3A**). **Fig. 3B** shows the induction of some representative genes of these processes, and **Fig. 3C** displays the gene-concept network of select differentially expressed genes associated with the indicated biological pathways including neutrophil degranulation, interleukin signaling, and HSF-1 activation.

**Figure 2.**
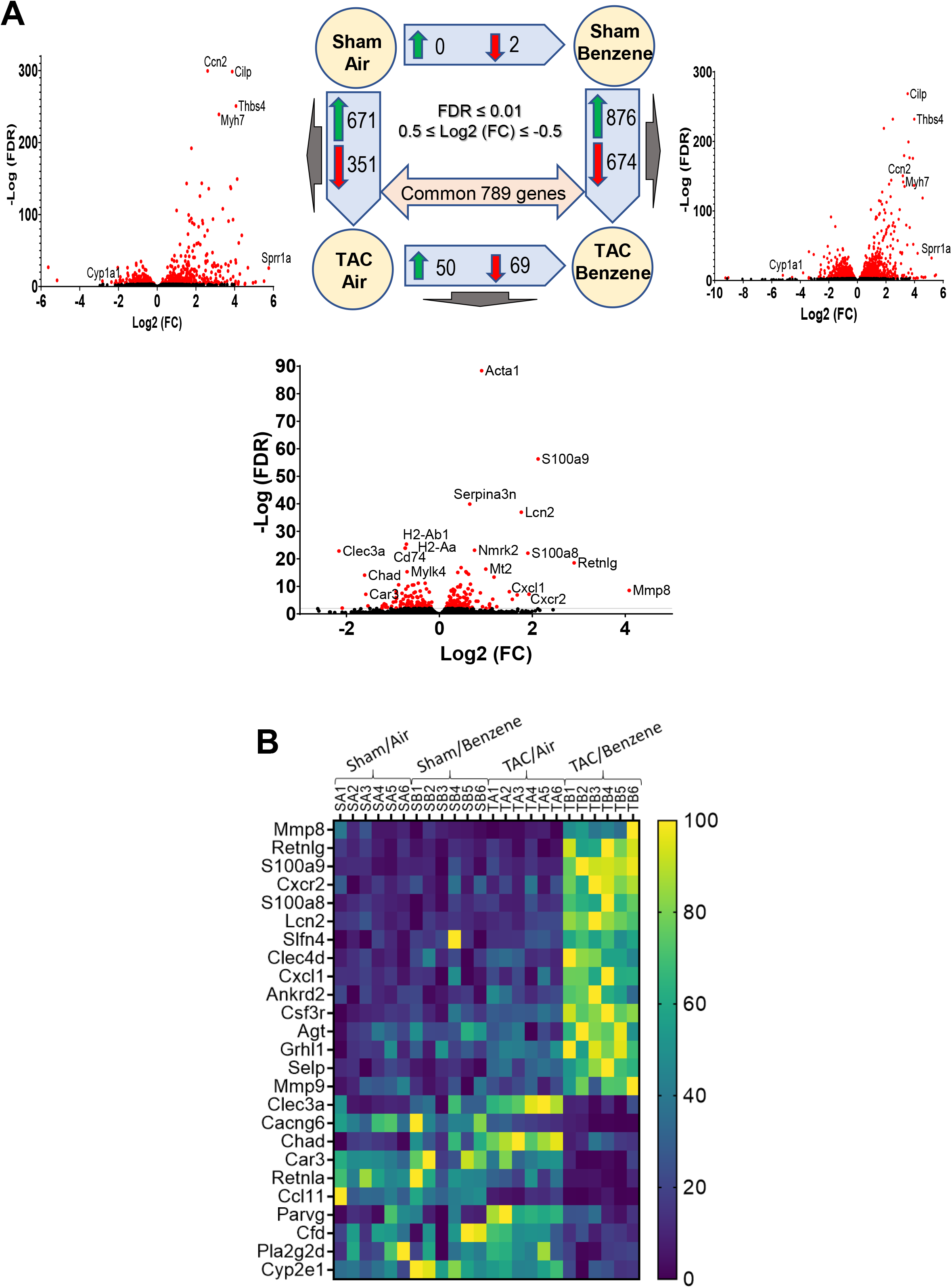
Differential expression of genes (DEG) in the hearts of TAC/Benzene-exposed mice. **A.** The number of differentially expressed genes (DEG) are depicted by corresponding arrows. The visual representation of DEG for three treatments are presented as volcano plots (FDR ≤ 0.01). **B.** Heatmap of the top 15 up-regulated and 10 down-regulated (TAC/Benzene vs. TAC/Air) protein-encoding genes in the four experimental groups.

**Figure 3.**
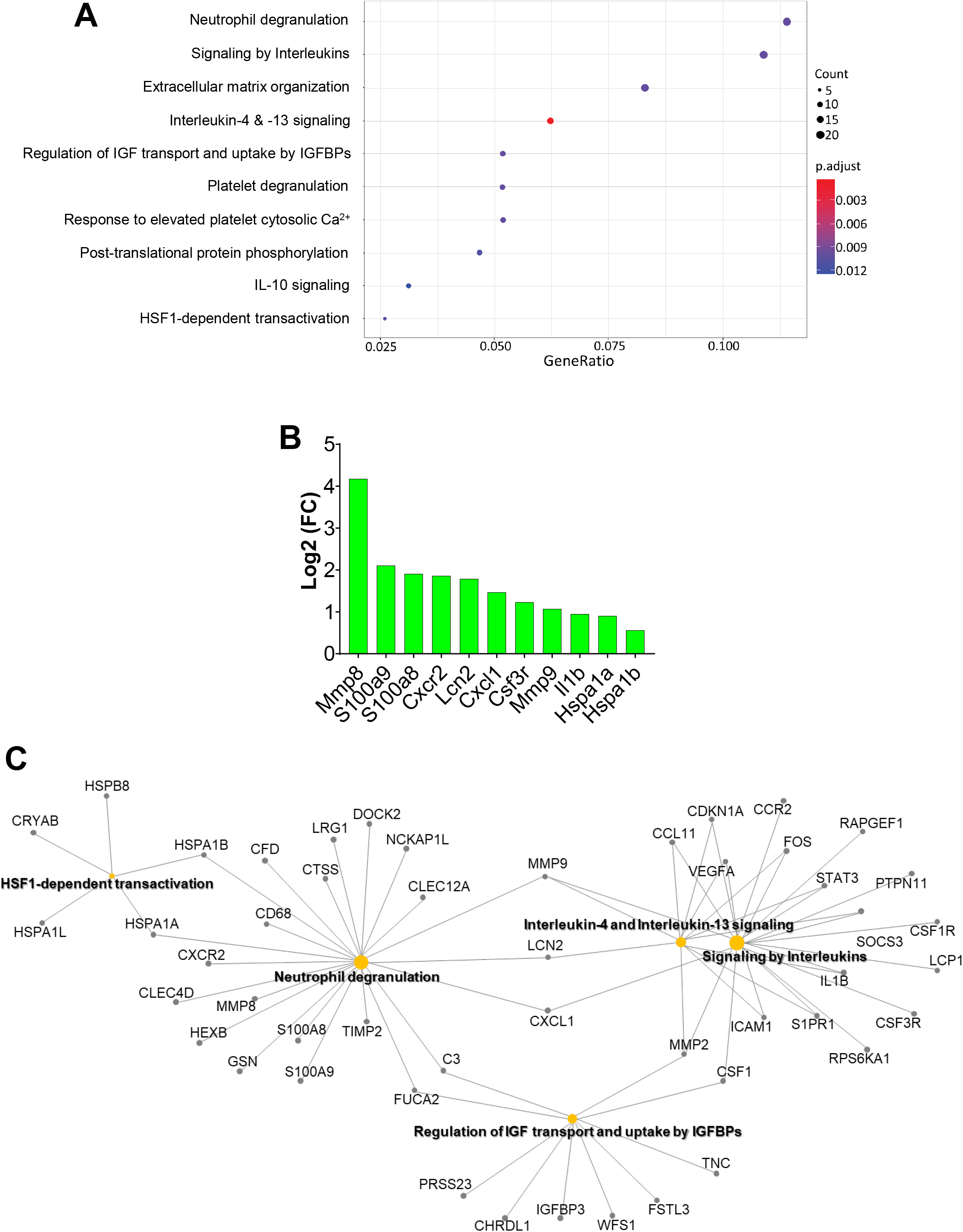
Pathway enrichment analysis of differentially expressed genes in the hearts of TAC/Benzene-exposed mice. **A.** Dot plot of enriched terms for TAC/Benzene vs TAC/Air DEG (FDR ≤ 0.01). IGF – Insulin-like Growth Factor, IGFBPs – Insulin-like Growth Factor Binding Proteins; **B.** The most upregulated and functionally important genes in TAC/Benzene vs TAC/Air groups. **C.** Gene-concept network depicting the linkage of genes and enriched biological pathways as a network.

### Benzene exposure augments TAC-induced neutrophil infiltration

Because our RNA-Seq data suggest that benzene exposure affects inflammatory response and neutrophil degranulation processes in TAC hearts, we examined the abundance of inflammatory cells in the TAC/benzene-exposed hearts. As shown in **Fig. 4A**, benzene exposure induced massive infiltration of S100A8-positive granulocytes in the hearts of TAC mice. These cells also co-stained with the pan-myeloid marker CD11b and myeloperoxidase, mostly expressed on neutrophils (**Fig. 4A and 4B**). Quantitative analysis shows >3-fold increase in S100A8^+^ granulocytes, CD11b^+^ myeloid cells and myeloperoxidase^+^ neutrophils in TAC/Benzene vs TAC/Air groups (**Fig. 4C**). The S100A8 staining did not co-localize with the markers of CD68^+^ macrophages and Thy1^+^ fibroblasts in TAC-Benzene hearts (***Supplemental* Fig. 2**). To examine the mechanisms by which benzene exposure augments neutrophil infiltration in TAC-instrumented hearts, we analyzed the RNA-seq data for the differential expression of adhesion molecules. As shown in **Fig. 4D**, benzene exposure did not affect the levels of adhesion molecules in the Sham-operated hearts, however, it significantly increased the transcription of Intracellular adhesion molecule-1 (*Icam1*), endothelial-selectin (*Sele*), and platelet-selectin (*Selp*) genes in the TAC-instrumented hearts. Expression of vascular cell adhesion molecule (*Vcam1*) in TAC-benzene hearts was comparable with the TAC/Air hearts. Together, these data indicate that chronic benzene exposure enhances the infiltration of neutrophils, possibly by activating adhesion molecules in the myocardial endothelial cells during pressure overload-induced cardiac dysfunction.

**Figure 4.**
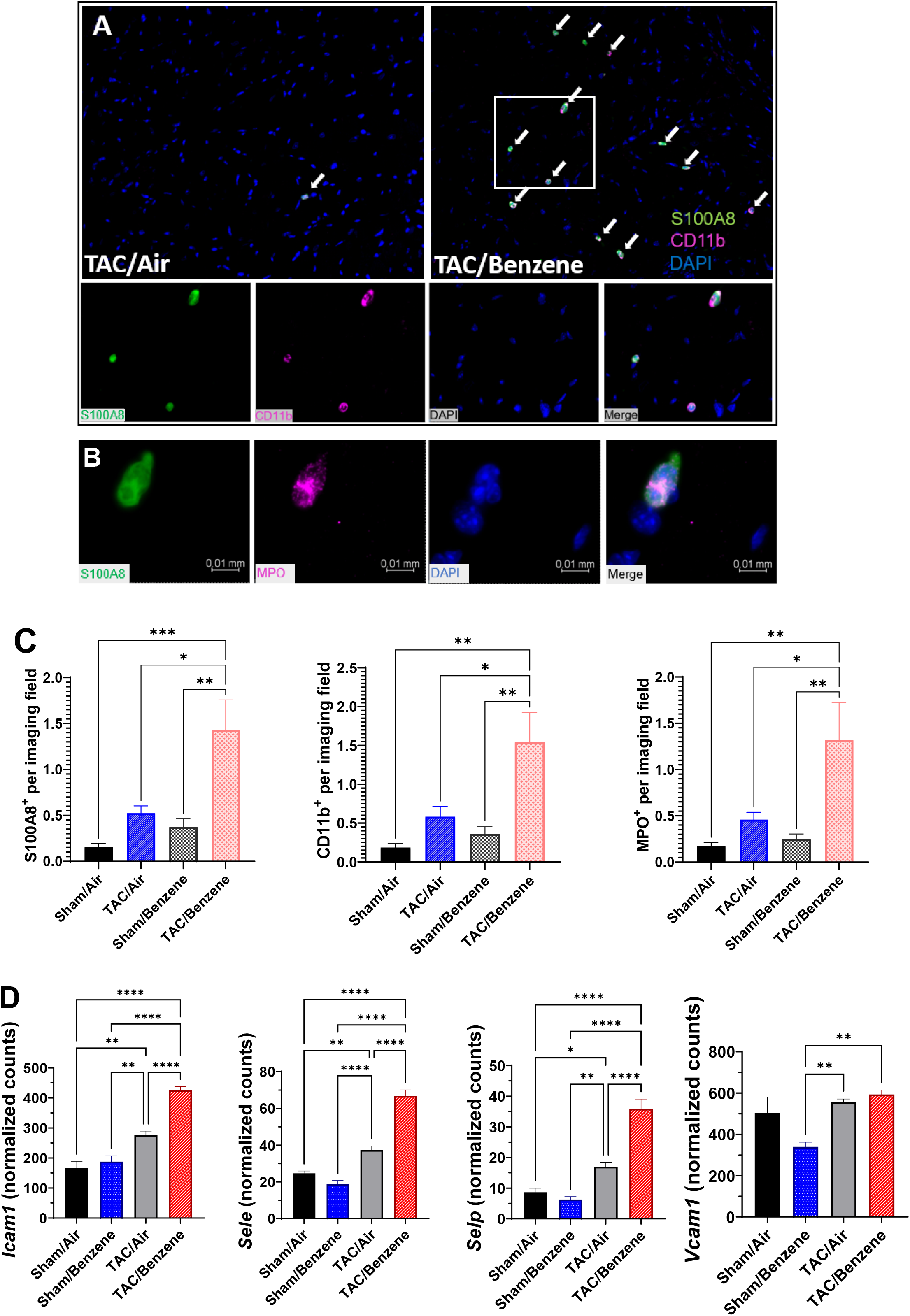
Immune cells infiltration into the heart of TAC/Benzene-exposed mice. **A.** Heart sections were stained for S100A8 (green) and CD11b (pink), nuclei (blue); **B.** Co-staining of S100A8 positive cells with myeloperoxidase (MPO) positive cells. **C.** Quantitative analysis of S100A8-, CD11b-, and MPO-positive cells. **D.** Expression levels of adhesion genes in the hearts of exposed mice. Values are mean ± SEM. *p<0.05, **p<0.01 and ***p<0.001 represent statistical significance between corresponding groups analyzed by One-way ANOVA. N=6/group.

### P-Selectin facilitates the adhesion of neutrophils adhesion to the cardiac microvascular endothelial cells

Complimentary *in vitro* experiments with CMVEC showed that benzene metabolites hydroquinone (5 μM, 24 h) and catechol (5 μM, 24 h) increased the transcription of P-selectin by 5-fold (**Fig. 5A**) and abundance of P-selectin protein by 1.5-2-fold (**Fig. 5B**). Incubation of CMVEC with catechol for 24 h augmented the adhesion of bone marrow-derived neutrophils to CMVEC (**Fig. 5C**), which was attenuated by pre-incubation of CMVEC with anti-P-selectin antibody (**Fig. 5D**). Hydroquinone and catechol also increased the expression of interleukin-8 (IL8, ***Supplemental* Fig. 3**), which can function both as a membrane-bound and soluble activator of neutrophil β2 integrin-mediated adhesion. Together, these observations suggest that benzene-induced neutrophil infiltration in the TAC hearts could be facilitated, at least in part, by the metabolic conversion of benzene to its active metabolites-hydroquinone and catechol.

**Figure 5.**
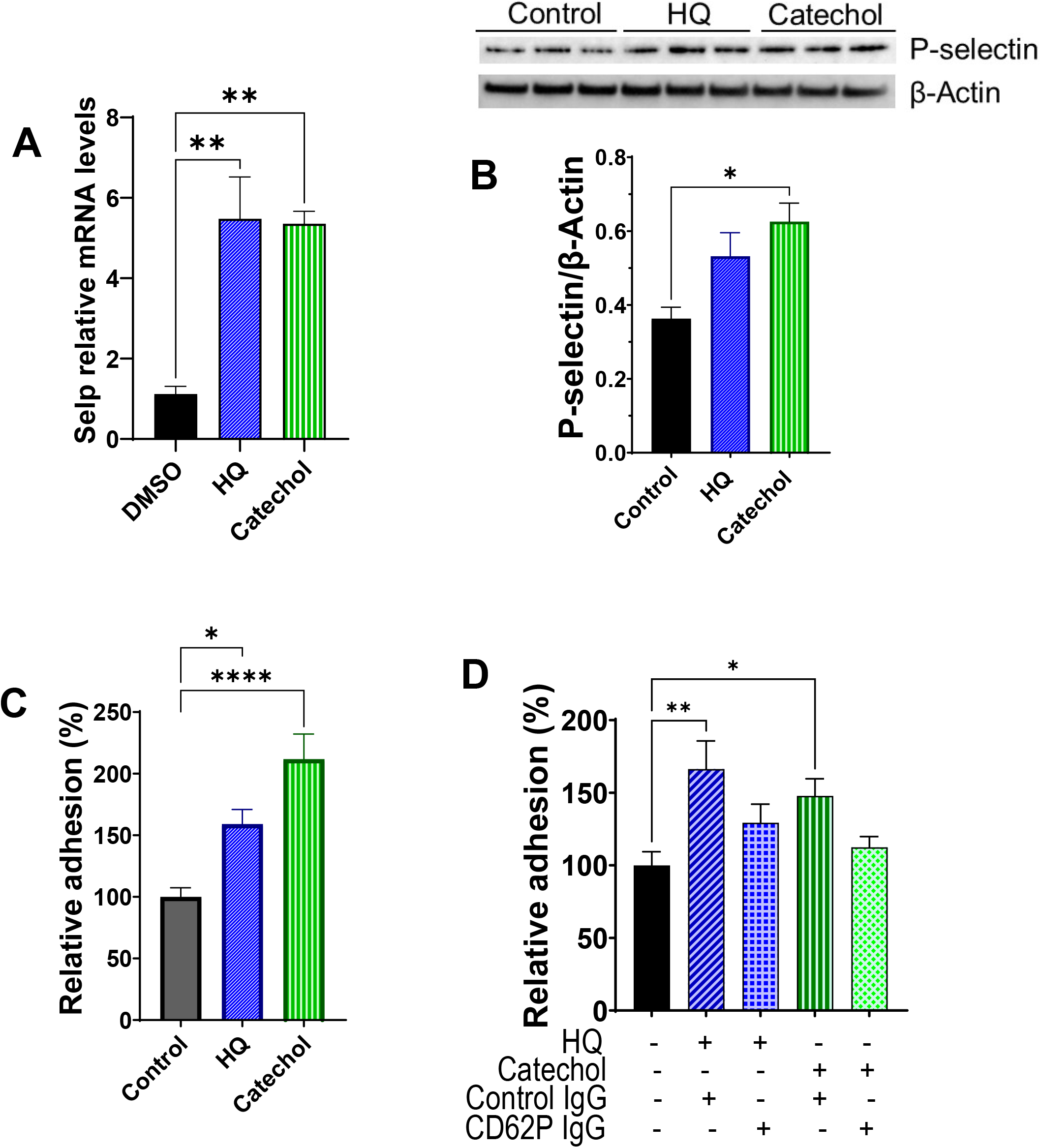
Benzene metabolites induce neutrophil adhesion to the cardiac microvascular endothelial cells. **A.** Transcription of P-selectin (*Selp*) in cardiac microvascular endothelial cells (CMVEC) treated with benzene metabolites hydroquinone (HQ; 5 μM, 6 h) and catechol (5 μM, 24 h). **B.** HQ (5 μM, 24 h)- and catechol (5 μM, 24 h)-induced upregulation of P-selectin protein in CMVEC. **C.** Adhesion of bone marrow-derived calcein-labeled murine neutrophils to CMVEC pretreated with HQ (5 μM) and Catechol (5 μM) for 24 hours. **D.** Inhibition of neutrophil adhesion to HQ (5 μM, 24 h)- and catechol (5 μM, 24 h)-stimulated CMVEC by anti-P-selectin antibody. Values are mean ± SEM. *p<0.05, **p<0.01, ***p<0.001, and ****p<0.0001 between corresponding groups analyzed by One-way ANOVA.

## DISCUSSION

The major finding of this study is that chronic benzene exposure exacerbates pressure overload-induced cardiac dysfunction. The TAC/Benzene-induced cardiac dysfunction is accompanied by endothelial activation and enhanced recruitment of granulocytes in the heart and differential regulation of genes involved in neutrophil degranulation, inflammatory signaling, and stress response. These studies corroborate recent reports that occupational and environmental exposure to benzene is associated with an increased risk for heart failure (25).

Inflammatory processes have long been implicated in the manifestation, progression, and consequences of ischemic heart disease (37, 38) and pressure- or volume overload-induced mechanical stress (17, 39–41). Pressure overload induces the expression of *ICAM1* (42) and consequent increase in the recruitment of neutrophils in the heart (39) and subsequent monocyte infiltration (17, 39). It diminishes TAC-induced cardiac hypertrophy and inflammation and sustains cardiac function (39), suggesting that neutrophils play a pivotal role in the manifestation of heart disease, especially heart failure. In mice, increased recruitment of S100A9 positive granulocytes, which co-stained with myeloperoxidase, suggests that TAC/Benzene exposure facilitates the infiltration of neutrophils into the hearts.

This is further supported by RNAseq data of cardiac tissue, demonstrating robust induction in the neutrophil specific *S100A8/A9, Mmp8, Lcn2*, and *Cxcr2* transcripts. S100A8/A9 is a member of alarmins or damage-associated molecular patterns (DAMPs) that are released by the failing heart (43–45) and subsequently activate resident macrophages, fibroblasts, and endothelial cells to secrete proinflammatory cytokines and adhesion molecules (46) and augment inflammation in the cardiac tissue (21, 23, 47). Systemic administration of S100A8/A9 facilitates the mobilization of neutrophils from the bone marrow (48) while local administration of S100A8/A9 leads to rapid recruitment of monocytes and neutrophils into the site of injury (49). Patients with myocardial infarction and elevated S100A8/A9 levels have poorer prognosis (50, 51). Overall, increased neutrophil infiltration in the hearts could play a pivotal role in benzene exposure-induced cardiac dysfunction in the pressure overload hearts.

Although precise mechanisms by which benzene exposure exacerbates pressure overload-induced cardiac dysfunction remain unknown, it is plausible that the observed enlargement of left ventricle dimensions in the hearts of TAC/Benzene mice could, at least in part, be due to the induction of neutrophil gelatinase-associated lipocalin (NGAL or Lcn2) whose expression is regulated by the S100A8/A9 complex (52). Lcn2 plays a central role in the ontogeny of cardiac hypertrophy and heart failure (53–55) and can bind Mmp9 to create a complex to stabilize Mmp9 protein and enhance metalloproteinase activity with its pro-angiogenic and pro-invasive properties (56, 57). Moreover, S100A8/A9 can diminish cardiac myocyte contractility, resulting in attenuated cardiac ejection fraction (58). Patients of chronic heart failure have elevated plasma myeloperoxidase levels (59), and pharmacological inhibition of myeloperoxidase diminishes leukocyte recruitment, and improves ejection fraction, end diastolic/systolic volume, and left ventricular hypertrophy following myocardial infarction (60). Therefore, it is conceivable that the augmented neutrophil infiltration with subsequent release of MPO and S100A8/A9 heterodimer from activated granulocytes might affect myocyte contractility and compromise cardiac function in TAC/Benzene mice.

Increased inflammation in the hearts of mice subjected to pressure overload followed by benzene exposure is further underscored by the increased transcription of chemokines, cytokines, and their receptors (e.g. Il1β, Cxcl1, and Cxcr2). Chemokines such as CXCL1 chemokine are well known to facilitate neutrophil migration into inflamed tissues (61), promote angiogenesis (62, 63) and regulate the recruitment of neutrophils and monocytes during cardiac remodeling and inflammation (64). Although the cellular sources (e.g., endothelial cells, granulocytes) of CXCL1 and other chemokines and cytokines in the hearts of TAC/benzene-exposed mice remain unknown, they likely augment neutrophil recruitment and migration. Benzene metabolites have been shown to increase the expression of cytokines and chemokines in the peripheral blood mononuclear cells (65) and endothelial cells (66). Our *in vitro* studies show that in murine cardiac microvascular endothelial cells, benzene metabolites hydroquinone and catechol increase the transcription of chemokine IL-8, which is known to bind to CXCR2 on neutrophils and trigger a signaling cascade that facilitates a firm monocyte adhesion with subsequent intravasation (67). Further studies are required to delineate the sources of chemokines and cytokines and the molecular mechanisms by which these molecules promote inflammatory signaling in the hearts of TAC/Benzene mice.

Induction of heat shock proteins (HSPs) such as *Hspa1a* and *Hspa1b* in Sham/Benzene hearts appears to be an adaptive response to circumvent inflammatory signaling. Induction of HSPs in response to chemical and mechanical stress enhances protein folding, inhibits inflammation and apoptosis, and provides cytoskeletal protection (68). While little is known about the role of HSPs in pressure overload-induced heart failure, heat shock factor 1 (HSF-1), the transcription factor that regulates HSPs is induced by TAC in rats and mice, and constitutive activation of HSF-1 ameliorates TAC-induced cardiac hypertrophy, fibrosis and cardiac dysfunction (69–71). Conversely, inhibition of HSF-1 impairs cardiac function in the pressure overload subjected hearts (69, 70). The HSF-1-mediated cardiac adaptation from pressure overload has been attributed, at least in part, to myocardial angiogenesis via suppression of p53 and subsequent induction of HSF-1 (72). HSF-1 can also improve endothelial function (73) and prevent inflammatory signaling (74). Although the contribution of the HSF-1/HSP axis in the etiology of cardiac dysfunction following TAC/Benzene exposure is unclear, it is likely that these pathways are activated to prevent benzene-induced endothelial activation. Further studies are required to examine the molecular mechanisms by which HSF-1-dependent transactivation prevents endothelial activation.

Collectively, our studies demonstrate that benzene exposure worsens pressure overload-induced cardiac dysfunction, which is accompanied by increased influx of neutrophils, alterations in cell-cell adhesion, and induction of genes associated with inflammation and stress response. These studies establish the plausibility that benzene exposure is sufficient to exacerbate TAC-induced cardiac dysfunction and provide insight about the underlying mechanisms.

## Acknowledgment

This study was supported in parts by NIH grants P42 ES023716, R01 HL149351, R01 HL137229, R01 HL146134, R01 HL156362, R01 HL138992, R01 HL122676, R21 ES033323, U54 HL120163, P30 GM127607, NIH S10 OD025178, and the Jewish Heritage Foundation grant OGMN190574L.

---

**Supplementary Table 1.** Echocardiographic parameters of transverse aortic constriction or sham operated mice followed by benzene or air exposure for 6 weeks.

| **Parameters** | **Sham/Air** | **Sham/Benzene** | **TAC/Air** | **TAC/Benzene** |
| --- | --- | --- | --- | --- |
| IVRT (ms) | 11.1 ± 0.6 | 12.6 ± 0.7 | 16.2 ± 2.0 | 17.8 ± 2.3* |
| LVAWd (mm) | 1.3 ± 0.1 | 1.2 ± 0.03 | 1.4 ± 0.04<sup>§</sup> | 1.4 ± 0.07 |
| LVAWs (mm) | 1.9 ± 0.07 | 1.8 ± 0.04 | 1.8 ± 0.06 | 1.7 ± 0.09 |
| LVPWd (mm) | 1.1 ± 0.04 | 1.0 ± 0.05 | 1.3 ± 0.1<sup>§</sup> | 1.2 ± 0.06 |
| LVPWs (mm) | 1.8 ± 0.06 | 1.5 ± 0.06 | 1.6 ± 0.09 | 1.5 ± 0.09* |
| HR (bpm) | 534 ± 11 | 514 ± 12 | 523 ± 9 | 549 ± 9 |
| Endo CO (mL/min) | 13.7 ± 1.01 | 15.6 ± 1.21 | 12.1 ± 0.98 | 11.2 ± 1.03<sup>§</sup> |
| SV (μL) | 25.9 ± 2.3 | 30.6 ± 2.7 | 23.1 ± 1.9 | 20.3 ± 1.7<sup>§</sup> |

Values are mean ± SEM. N=10-12 per group. \*p ≤ 0.05 vs Sham/Air, <sup>§</sup>p ≤ 0.05 vs Sham/Benzene. Statistical analyses were performed by One-way ANOVA.

**Supplementary Figure 1.**
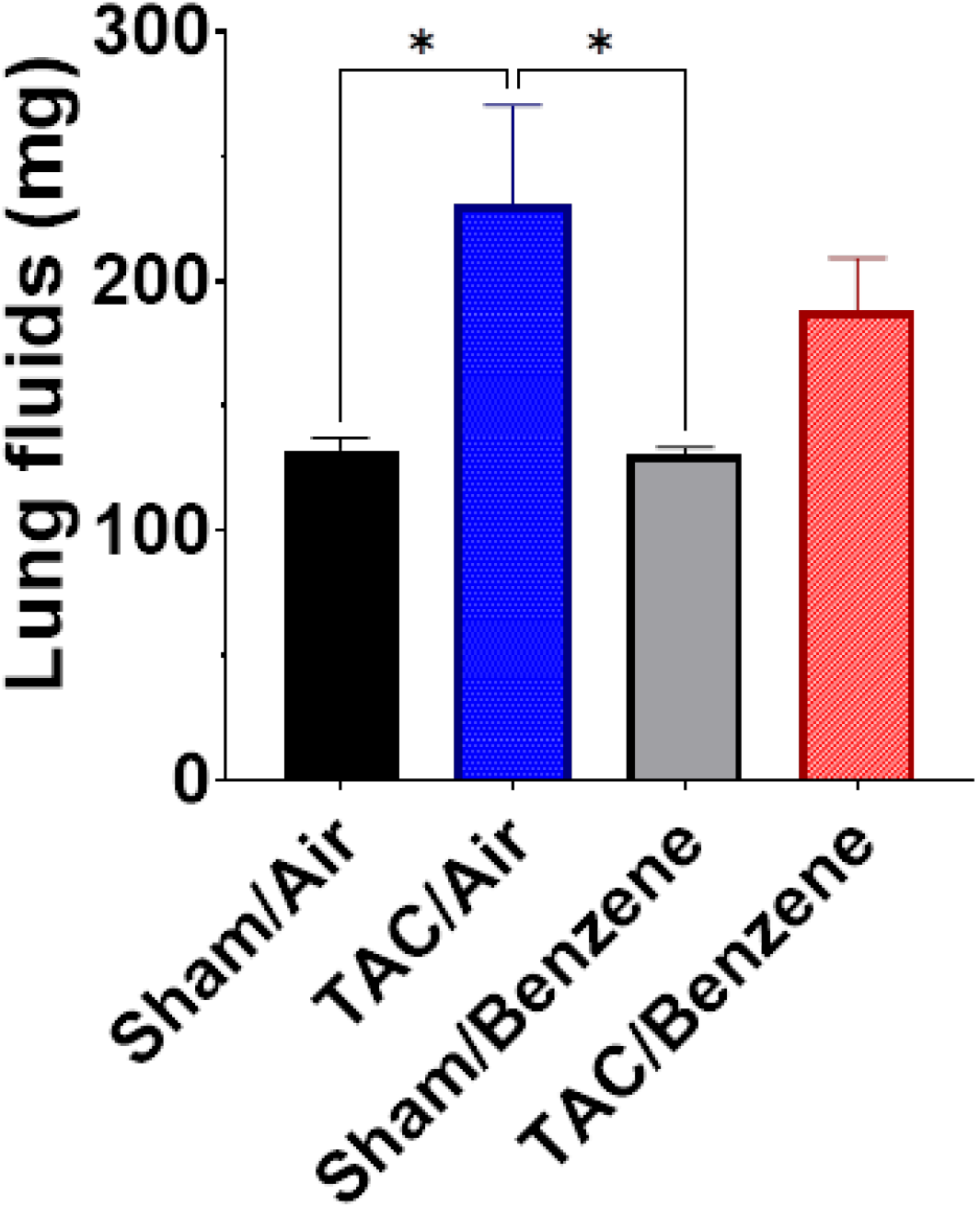
Pulmonary edema in transverse aortic constriction or Sham operated mice exposed to benzene or air for 6 weeks. Accumulation of the fluid in the lung was measured by subtracting the dry weight of the tissue from the wet weight. Values are mean ± SEM, N=10-12 per group. *p<0.05 between corresponding groups. Statistical analyses were performed by One-way ANOVA.

**Supplementary Figure 2.**
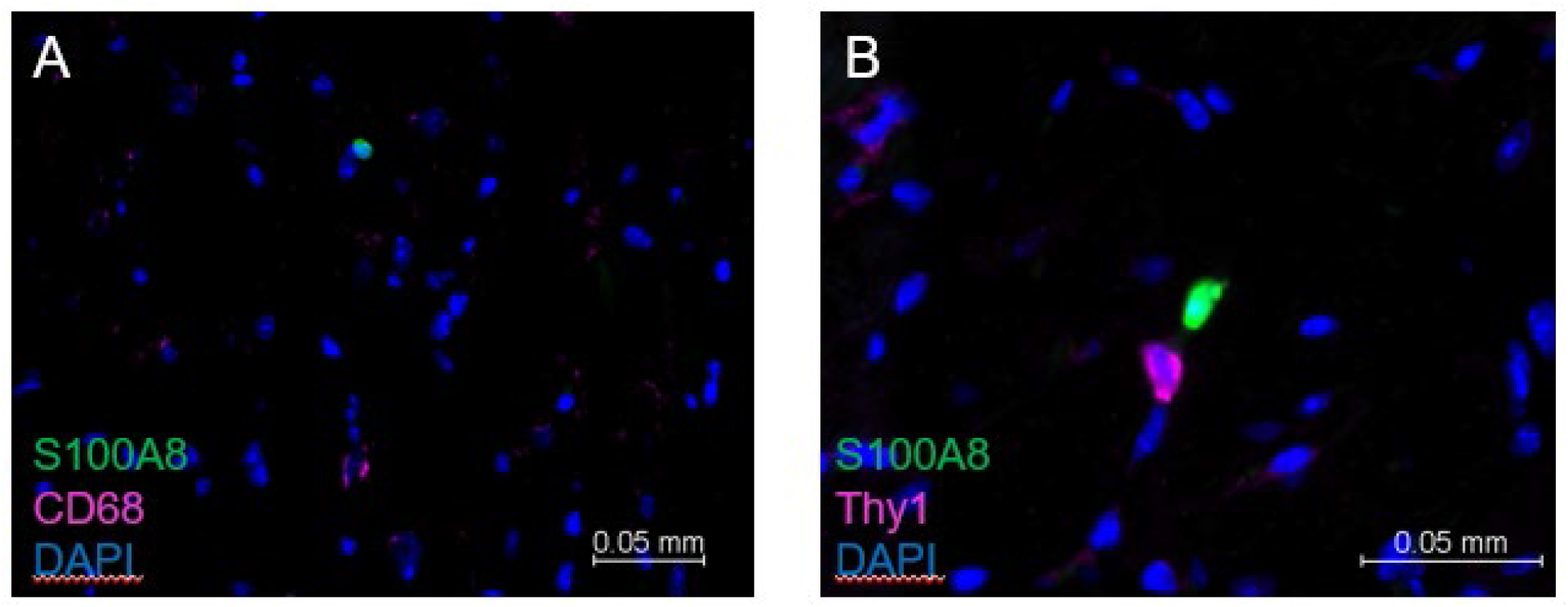
Staining of immune cells in the hearts of TAC/Benzene-exposed mice. Hearts from the TAC/Benzene mice were stained with granulocyte marker S100A8, macrophage marker CD68, or fibroblast marker Thy1. Nuclei were stained with DAPI (blue).

**Supplementary Figure 3.**
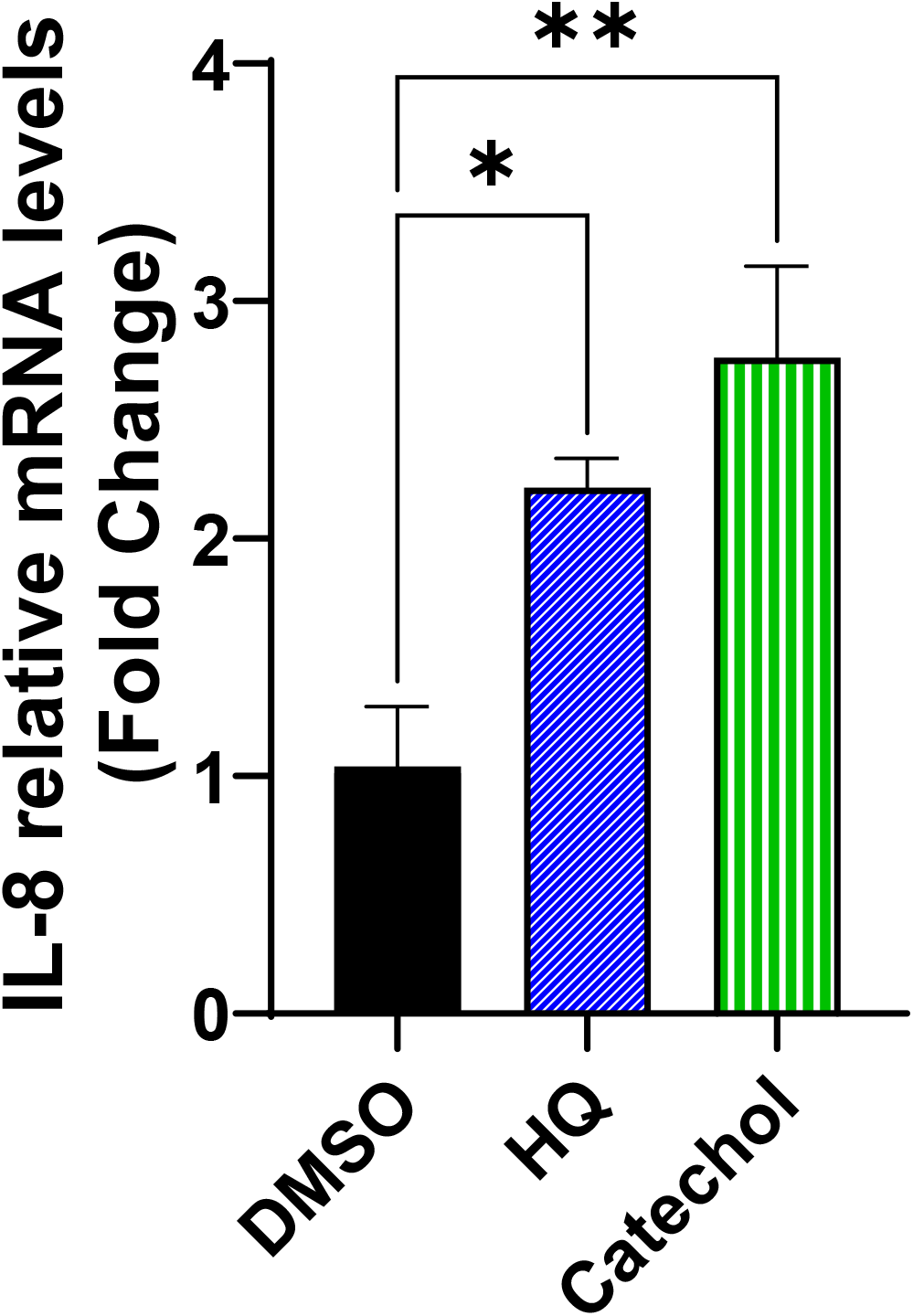
Expression of chemokine IL8 in hydroquinone (HQ, 5 μM/24h) and catechol cardiac (5 μM/24h)-treated cardiac microvascular endothelial cells. Values are mean ± SEM, N=3 per group. *p<0.05, **p<0.01 between corresponding groups. Statistical analyses were performed by One-way ANOVA.

